# Cause for pause before leaping to conclusions about stepping

**DOI:** 10.1101/085886

**Authors:** Ariel Zylberberg, Michael N. Shadlen

**Affiliations:** Howard Hughes Medical Institute, Kavli Institute and Department of Neuroscience, Zuckerman Mind Brain Behavior Institute, Columbia University, New York, NY 10032, USA

## Abstract

Many neurons in parietal and prefrontal association cortex undergo gradual changes in firing rate during the formation of some perceptual decisions. These dynamics are often ramp-like increases or decreases depending on the sign and strength of the sensory evidence and are thus hypothesized to represent the accumulation of noisy samples of evidence, analogous to biased diffusion. This idea was challenged recently. An analysis of sequences of action potentials recorded from neurons in the lateral intraparietal cortex (area LIP) suggests that the spikes on single trials are explained by rates that undergo a discrete step from an intermediate rate to either a low or high rate at a random time during deliberation. The average of such steps, like the average of biased diffusion, is consistent with the ramp-like firing rates observed in LIP, but a Bayesian model comparison deemed stepping superior. Here we show that a shortcoming in the mathematical depiction of drift-diffusion led to a severe bias in the model comparison. We conclude that at present there is no compelling evidence that favors the stepping account.

## Introduction and Results

Many decisions benefit from the accumulation of noisy evidence, acquired sequentially over time. This accumulation, termed a decision variable (DV) comprises a deterministic component—conceptually, the integrated expectation of the samples—and the accumulation of unbiased noise. Thus the process is likened to diffusion plus drift, random walk plus forced march, a biased random walk, or Brownian motion plus drift. The process is ubiquitous in biology and chemistry—hence the variety of terminologies. In psychology, it coordinates error rates with the speed of decisions in free-response tasks. The decision terminates when the DV reaches a bound or threshold, which simultaneously resolves the decision that is made and the time taken to make it. Empirical support for these models comes from fits to human and animal choice behavior in a wide variety of tasks, including perceptual, economic and social decisions, memory retrieval and numerical comparisons [1-6].

A neural correlate of this evidence accumulation process has been characterized in cortical and sub-cortical association regions of the brain while animals make difficult perceptual decisions. In macaques, many studies focused on a region of the parietal cortex referred to as the lateral intraparietal area (LIP), and, in particular, in a subset of neurons which exhibit persistent activity during memory-saccade tasks [7]. When a visual stimulus instructs a monkey to plan an eye movement to a particular location in space (the neuron’s response field, RF), these neurons rapidly increase the firing rate and maintain an enhanced response until the monkey is cued to execute the saccade. When the saccadic direction is instructed with an ambiguous cue, such as the direction of motion of a cloud of dynamic random dots, the same neurons exhibit more gradual changes in firing rate. The firing rate increases when the evidence and choice favor the target in the RF (T_in_), and the rate decreases when the evidence and choice favor the opposite target (T_out_). Thus the trial-averaged rates appear ramp-like with a rate of rise or decline that reflects the strength and direction of the motion evidence. Regardless of task difficulty, the firing rates reach a common level before the eye movement, consistent with a threshold or bound applied to the neural representation of accumulated evidence.

The observation of ramp-like responses of LIP neurons during the deliberation phase of a decision raises the question, what do these averages comprise? In other words, what are the response dynamics on single trials? The answer that is most consistent with choice and reaction times is that the rates on individual trials resemble 1-dimensional Brownian motion plus deterministic drift, represented by the discrete stochastic difference equation

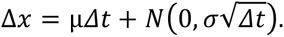

The latent rate, *x*(*t*), is the cumulative sum of samples from a Normal distribution with mean μΔt and variance *σ*^2^Δt, here split into a deterministic first term and a stochastic, Brownian term. It is often represented in continuous time by

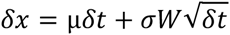

where *W* is the Wiener process. We refer to *x*(*t*) as a latent rate because it is not directly observed but instead manifests as a sequence of spikes.

To resolve the spike rate of a neuron in time, neurophysiologists typically repeat trials and average the spike occurrences in narrow time bins. This reveals the deterministic component of *x*(*t*), the ramp, but it suppresses the stochastic component, which is the hallmark of Brownian (diffusion) dynamics. However, it is possible to observe the signature of this process in 2^nd^ order statistics—the evolution of both the variance of *x*(*t*) (e.g., a linear increase as a function of time) across trials and the autocorrelation of *x*(*t*) within trials (a function of both the lag between samples and the time of the first sample) [8, 9]. Thus it has been surmised that the ramp-like averaged firing rates from neurons in LIP comprise samples of latent rates on individual trials that conform to a drift-diffusion like process.

A recent study challenged this view ([10]; hereafter referred to as *the study*), concluding instead that the gradual change in firing rate observed during the motion discrimination task may be explained as an average over trials in which the activity undergoes steps from a neutral to a low or high firing rate. If such steps were asynchronous across trials, the average could approximate a ramp, which would be indistinguishable from a ramp composed of the average of drift-diffusion processes. Using data from Alex Huk’s lab [11], Latimer et al. conducted a Bayesian model comparison of ‘stepping’ and ‘diffusion’ to determine which model best accounts for the response of single neurons recorded from LIP while monkeys performed a random-dot motion discrimination task. The comparison favored stepping over diffusion in ~3/4 of the neurons. Importantly, because the models are fit to the spiking activity on single trials, they are in principle capable of distinguishing between stepping and diffusion accounts of the latent rates.

In a brief Technical Comment, we pointed out numerous deficiencies in the data set and model comparison, which question the legitimacy of the conclusions [12]. We encourage the reader to examine the full Comment and the authors’ reply [13]. Here we focus on one important shortcoming, concerning the negative latent firing rates. Owing to its technical nature, this issue was mentioned only briefly in the Technical Comment and ignored in the reply. In what follows, we explain why the study’s characterization of diffusion led to severe bias in the model comparison. Put simply, their diffusion model cannot account for late spikes in experimental data when the evidence would tend to drive the neurons to decrease their firing rates. We first provide an intuition for the particular implementation of diffusion in *the study* and point out its limitation. We then explain why, despite this limitation, the model appeared to pass a simple validation test. We then show how the implementation biased the analysis of the data from the Huk lab, as well as the data from an earlier reaction time experiment [14] that was analyzed in supplementary material of *the study.*

In the diffusion model, as implemented in *the study,* the spike rate on any one trial conforms to a random walk plus a constant linear *drift,* which depends on the strength and direction of motion. The rate is frozen at an upper bound if it is reached, whereas there is no corresponding lower bound that would freeze the firing rate at a low value. Until the upper bound is reached or until the trial ends, the random walk can achieve arbitrarily large negative values, which are “soft rectified” to zero (approximately). When the drift is negative, most trials do not terminate in the upper bound. If the model assigns a negative drift rate, it must also predict very few spikes late in the trial. For example, if the data were to exhibit a decline in firing rate early in the trial, say in response to strong motion in the direction opposite to the one favored by the neuron, the model cannot capture this with a negative drift rate without incurring severe penalty for spikes occurring later in the trial. Crucially, this does not constitute a general limitation of diffusion models but of the specific instantiation that the authors chose to implement.

To understand why this feature favors stepping over drift-diffusion, it may be helpful to consider a toy example. Consider two trials from an experiment in which strong motion favors the choice target in or out of the neural response field (T_in_ and T_out_, respectively). These are shown by smooth firing rate functions in Figure 1A. For the T_in_ trial (blue trace), the response is fixed to the firing rate determined by a threshold or bound. For the T_out_ trial (red trace), the response is fixed at a value achieved when another competing set of neurons—whose RF aligns to that target—reach a positive threshold. Importantly, they are also frozen to some value for the remainder of the trial, but this will not be stereotyped because the stopping is controlled by a threshold operation on a different (positive going) process. This is the pattern that one sees in data from many experiments (e.g., [15, 16]). In the example, when the diffusion terminates the firing rate of the neuron is close to 10 spikes per second (sp/s), similar to the average T_out_ rate in *the study* (see their Figure 4, adapted below as Figure 4).

**Figure 1.**
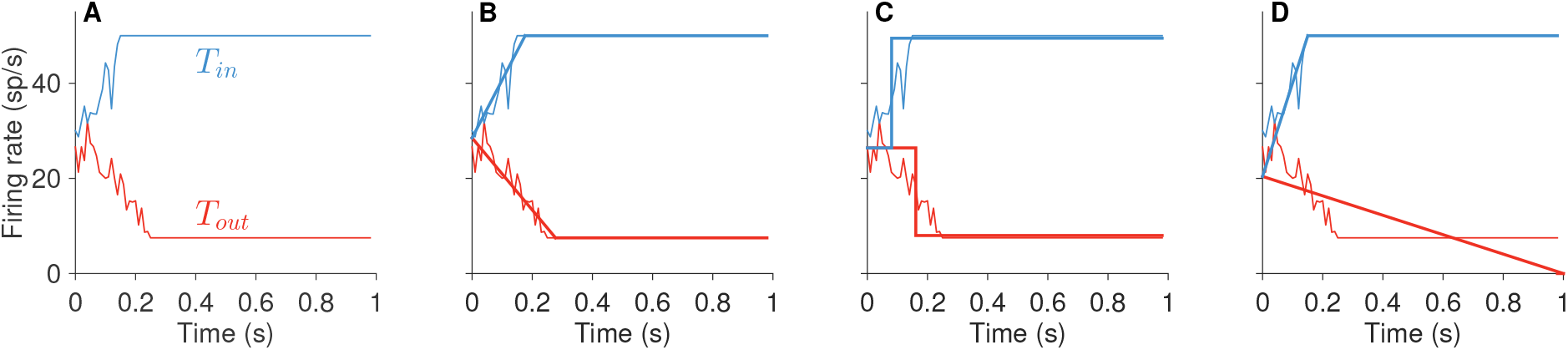
Fits of steps and ramps to hypothetical rates. Hypothetical firing rate trajectories (**A**) and their fits by a ramping model (**B**), a stepping model (**C**), and a ramping model with unstopped negative rates (**D**).

Now, consider what would happen if we attempt to fit a set of ramps—that is, the deterministic component of drift-diffusion—or steps to these traces. If we allow the ramp to freeze at an upper or lower termination rate, the model captures both the gradual change and the later constant response in the simulated rates (Fig. 1B). The stepping model does slightly worse (Fig. 1C). It incurs error because it cannot explain the intermediate rates, but the ramps in the example are brief, and the error is minimized by setting the time of the step near the midpoint of the ramps. A ramp that is not allowed to freeze at a lower termination rate would produce a fit more like the one in Figure 1D–a clear miss for the T_out_ trial. To minimize error, the slope of the ramp is set to a value that is less negative than it ought to be.

The exercise provides an intuition for why a model that does not allow the negative-going firing rates to plateau at non-zero values may lead to a biased comparison between the diffusion and stepping models. In what follows we show that the diffusion model in *the study* lacks a mechanism to stop negative going rates, just as in Fig. 1D; then we use the algorithms supplied by the Pillow lab (https://github.com/pillowlab/StepRampMCMC) to fit neural responses and simulated data.

The lack of a mechanism to stop negative going latent rates in *the study* is clear from the equations that describe the dynamics of the diffusion model, which we reproduce here from section 2.1 of their supplementary materials. The parameters of the model are represented by the set *Φ =* {*β*_1:*c*_, *x*_0_, *ω*^2^, *γ*}, where *β*_1:*c*_ are the drift terms, one value per coherence, *c; ω2* is the diffusion term, which determines the amplitude of the Gaussian noise added to the process at each time step; *x*_0_ is value of the latent rate at the start of the process; *γ* determines the height of the upper bound. The full model is specified by these equations:

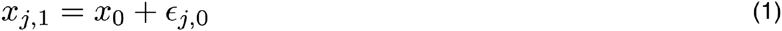

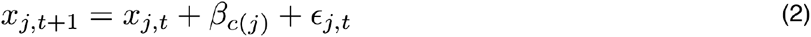

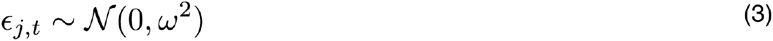

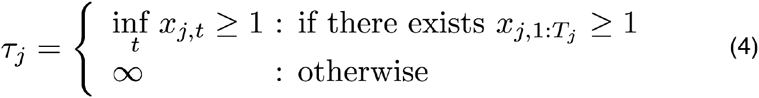

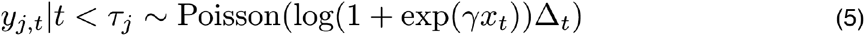

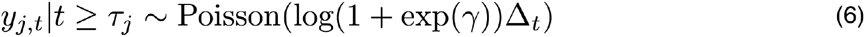

Equations 1-3 describe the evolution of the drift-diffusion process, where *x*_j,t_ is the state of the diffusing particle for trial *j* at time-step *t*, and ɛ is a sample of Gaussian noise. In equation 4, τ is the time at which an upper bound (at *x =* 1) is first hit (or ∞ if the upper bound was not hit). Equations 5-6 specify the conversion from the drift-diffusion process *x* to the spike counts *y.* The spike counts are samples from a Poisson distribution. For times lower than the bound-crossing time τ (equation 5), the parameter of the Poisson distribution depends on *x,* scaled by the bound height *γ.* For times greater than τ, the spike counts are independent of *x* (equation 6). Notice that while the diffusion process does not stop at the upper bound, it is equivalent to a model that implements this feature, because the spike counts do not depend on the value of *x* after the bound-hitting time. The firing rates are fixed at a value just greater than *γ* (equation 6). Unless *x* has reached 1, however, the process is unbounded. Equation 5 ensures that negative values of *x* are soft rectified to spike rates just above zero spikes per second.

Figure 2 illustrates the drift-diffusion process *x* (Fig. 2A) and the spike rates (Fig. 2B) that are obtained from simulating this process. The figure shows trials of three different motion coherences: negative high (in red), zero (black), and positive high (blue). When diffusion paths go negative, the soft rectification function forces the firing rates to be close to zero. If the latent rate reaches the bound, it is frozen at the value of the bound (*γ*) for the rest of the motion viewing epoch. There is no lower bound in the model which would freeze the rate at some minimum. Instead the diffusion process can drift to large negative values, which the soft rectification converts to a value close to zero—the logarithm of (1 + *e* raised to a negative number). This means that if a negative drift rate is needed to explain a fast drop in firing rate for trials with strong motion away from the neuron’s RF, the process should generate very few spikes in the latter part of the motion viewing epoch. For strong motion, this can be most of the trial epoch under consideration.

**Figure 2.**
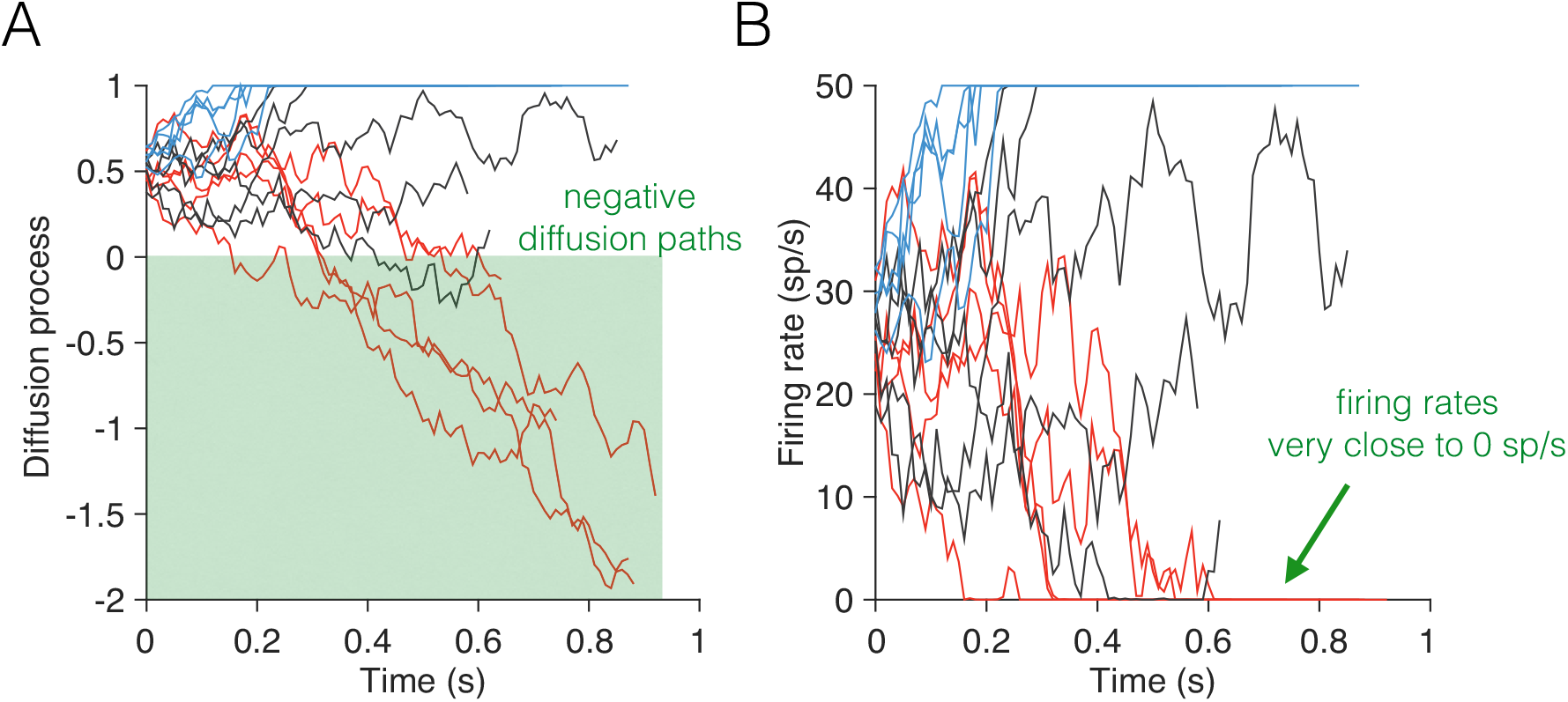
Example diffusion paths and latent firing rates for the diffusion model of *the study*. The motion strength is determined by the drift coefficient *β*, which is set to -0.02, 0, and 0.02 for the red, black and blue traces respectively. **A,** Sample diffusion processes *x* (five traces per motion strength). Negative-going processes are never stopped and continue the diffusion even when *x* becomes negative. **B**, Firing rates obtained after scaling and soft-rectifying the diffusion paths in A. Notice that the negative-going diffusion process are converted by the soft-rectification function into firing rates of ~0 spikes/second.

We used the code supplied by the Pillow lab to show that negative latent rates led to mistaken validation of the diffusion model. We compared two sets of simulations. The first recapitulates the simulations of diffusion used to validate the diffusion model in *the study.* It consists of simulating spikes under the equations above, for 5 coherence levels and 20 trials per coherence. Examples of latent firing rates and the sampled spikes are plotted in Figure 3A. Notice the near-zero spike rates for the negative coherences (red traces) and the near absence of spikes in the example rasters late in these trials. For the second simulation (Fig. 3B) we simply added a constant offset of 10 sp/s to the firing rates before using them to sample the spikes. Notice the presence of occasional spikes late in the trials for the negative coherences (red tics, Fig. 3B).

**Figure 3.**
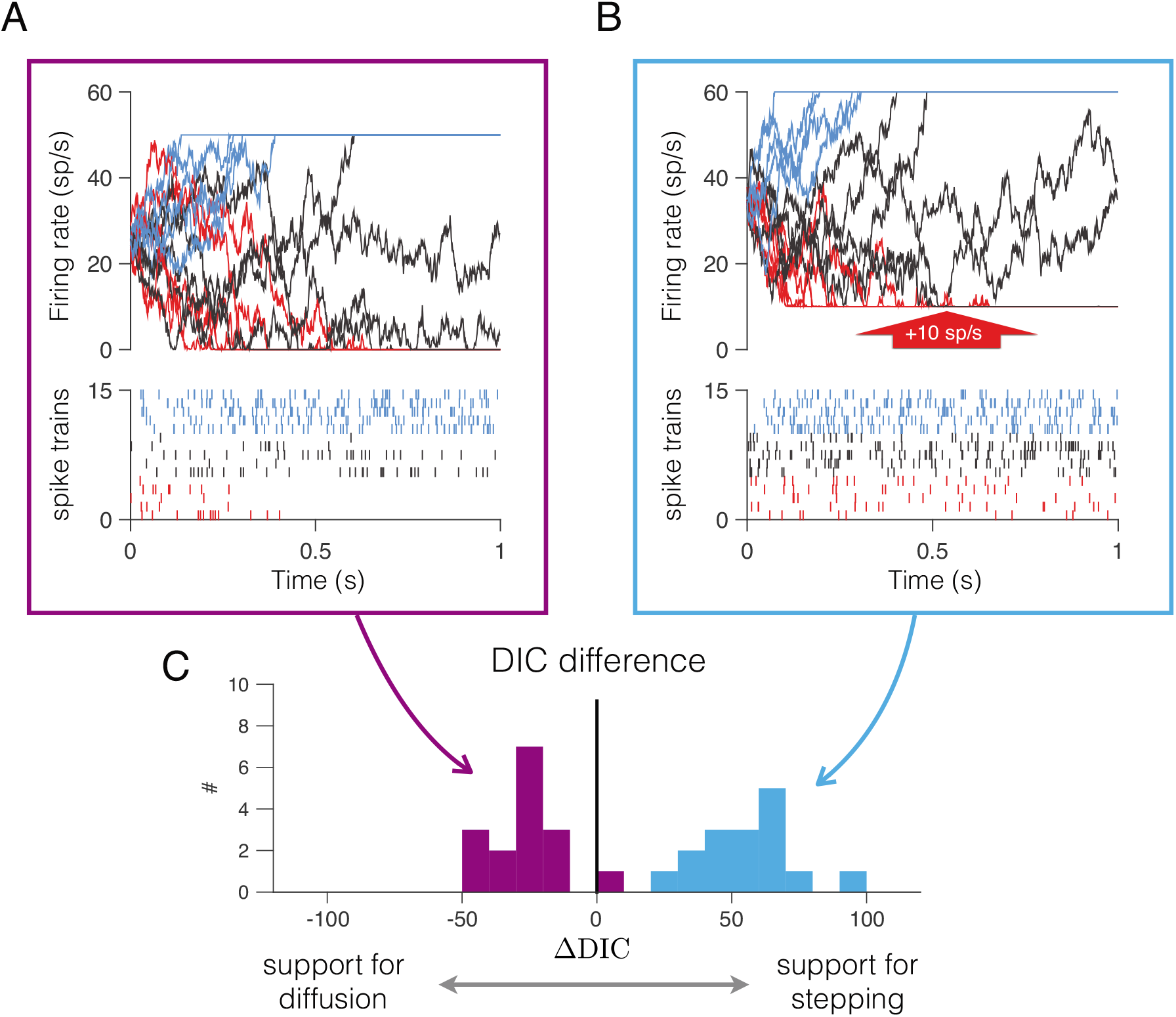
Failure to validate the diffusion model. **A**, Example trials simulated using the diffusion model in *the study.* Firing rates (top) are the soft rectified latent rates, corresponding to three motion strengths: strong T_in_ (blue), 0% coherence (black), and strong T_out_ (red). Each rate gives rise to sequence of spikes (bottom) via a nonstationary Poisson point process. **B**, Same simulated rates as in panel A plus a constant offset of 10 sp/s. Notice the occurrence of late spikes in the strong T_out_ condition. **C**, Model comparison. Stepping and diffusion models were fit to simulated data using the models in panels A and B (5 motion strengths; 20 trials per motion strength). Each of the colored histograms show the distribution of the ΔDIC statistic from fits to 32 simulated data sets. Each data set is fit with the stepping and diffusion models using the Pillow lab code. Negative ΔDIC supports diffusion; positive supports stepping. The purple histogram replicates the validation of the model: support for diffusion in 31 cases. The blue histogram shows strong support of stepping. Both sets of data were generated using diffusion. In the simulations, trial length were sampled uniformly from the range 0.5 and 1 s. Model parameters were: *β*_1:*c*_={-0.02,-0.01,0,0.01,0.02}; *x*_0_=0.5; *ω*^2^=0.005; *γ*=50.

Because both simulations were generated with diffusion, a comprehensive model comparison should favor diffusion over stepping in both cases, but that is not what we observed. We compared diffusion and stepping using the difference in Deviance Information Criteria (ΔDIC), as in *the study.* A positive ΔDIC indicates support for stepping; negative for diffusion. Figure 3C shows histograms of ΔDIC for 32 simulations (20 trials per 5 coherence levels per 16 simulations of both types). For the simulations like those in Figure 3A, the statistic supports diffusion, with one exception (purple histogram). For the simulations with the 10 sp/s offset, the statistic supports stepping and does so strongly. It misidentifies diffusion for stepping merely because of the late spikes, which are common in neural data but not in “validation data” generated from a model with soft-rectified negative latent rates. The exercise provides a cautionary tale about model validation, namely that it is only as good as the generative model that it seeks to validate. In this case, since LIP neurons typically exhibit late spikes even when the monkey chooses T_out_, it leads unsuspecting readers to confuse an inadequate depiction of diffusion as support for stepping.

The problem with the parameterization of the diffusion model is also apparent in the fits to neural data. Figure 4 reproduces the relevant panels from the fit of the diffusion model to the data from the Huk lab. It shows the same type of pathology illustrated in our toy example (Fig. 1). The upper left panel shows the PSTHs (averaged over the 40 cells) calculated from the spike counts in 25 ms windows. Colors indicate the motion strength and the direction of motion (T_in_ in blue, T_out_ in red). The mean firing rate on trials with strong motion in favor of T_out_ (labeled “-high”, Fig. 4) decays rapidly and then plateaus at a rate of ~10 sp/s (see the red trace in the upper-left quadrant of Fig. 4). The plateau is similar to that observed in previous studies (e.g., Figure 8 of [15]). The upper right panel shows the mean rate derived from the diffusion-model fits to the spike responses on each trial. Because the diffusion model can only reach a plateau at the firing rate of the upper bound, or at a firing rate close to 0 sp/s, it cannot reproduce the plateau observed in the data. Instead, the model settles to an almost linear decay in firing rate, like the red trace in the upper-right quadrant of Figure 4.

**Figure 4.**
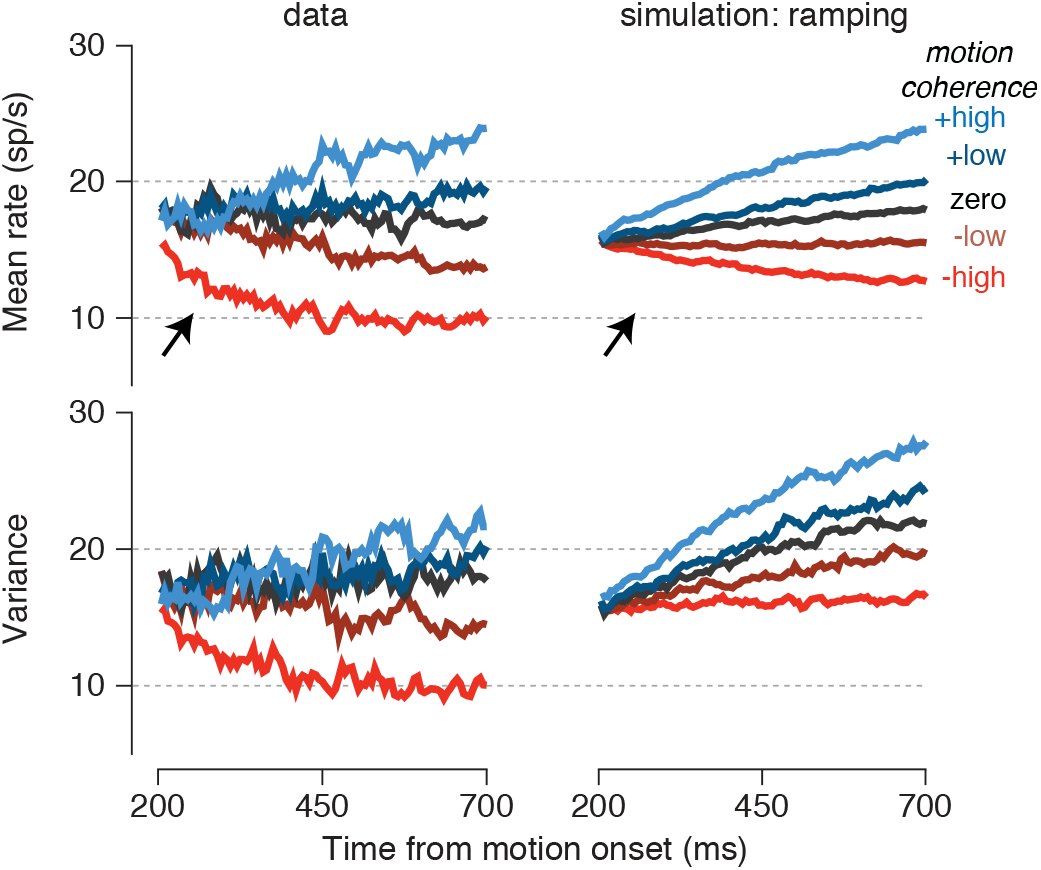
Unbounded negative latent rates bias the diffusion fits. For the negative going rates (labeled -low and -high), the diffusion model fails to capture the negative drift rate that is evident in the left part of the traces (arrows), corresponding to the epoch of evidence accumulation. The model assigns a shallower negative-going rate (top right) than is evidence in the early epoch of the data. It also mixes in positive going rates on some trials to achieve an average that plateaus above 10 sp/s. This mixture has consequences for the predicted variance of the firing rates (bottom right), which is clearly at odds with the variance in the neural data (bottom left). Adapted from Figure 4 of Latimer et al.

Unstopped negative rates appear to explain another peculiar feature of the diffusion fits in *the study,* shown on the bottom of Figure 4. Even without a mechanism that stops the negative-going rates, the plateau at positive firing rates can be approximated by mixing trials of low and high firing rate, as occurs when the drift rate is close to zero. For instance, from simulations of the diffusion model (with the parameters of Table S2 of *the study),* we observed that even for strong negative motion there is a considerable fraction of the trials (>20%) that reach the upper bound, and therefore the average will seem to plateau at a nonzero rate, even if no single latent rate does so. This solution does not save the model from a bias. It merely replaces the low likelihood of observing a late spike, given a latent rate near zero, with another low likelihood, given an inappropriately high latent rate.

Further, it predicts that for the diffusion model, the spikes occurring late in the trial will result from a mixture of two states, one with very low firing rate and another with a firing rate at the upper bound. This in turn predicts that the variance of the spike counts increases as a function of time even when the firing rates decrease with time. This is indeed what was observed from the fits of the diffusion model. The lower-right quadrant of Figure 4 shows the variance of the spike counts computed over the same 25 ms windows as the average rates and averaged across neurons (see section 2.4 of Supplementary Materials of *the study*). The population variance increases with time even when the rates do not (‘-high’ and ‘-low’ traces in the right panels of Figure 4). In contrast, the observed variance in data tends to parallel the mean, decreasing for T_out_ motion (lower-left quadrant of Fig. 4). While *the study* interprets this result as evidence against the diffusion model and in favor of stepping, we interpret it as a result of poor parameterization of the diffusion model, which does not allow the negative going rates to plateau at non-zero values. Again, it does not reflect a general limitation of the drift-diffusion account, but a limitation of the specific implementation of the diffusion process in *the study.*

Of course, the stepping model is well equipped to explain a nonzero firing rate plateau associated with strong motion in the “negative” direction. It sets the parameter of the downward jump to match this (*α*_*out*_ in Table S1 from *the study*). The variation in this parameter permits a test of our claim that the late spikes in the negative rates are a source of bias in the model comparison. We predicted that the bias should be reduced for cells with low *α*_*out*_. Indeed, of the 40 cells analyzed in *the study*, 9 exhibited *α*_*out*_ less than 2 sp/s (Table S1 from *the paper*), and of these, 5 were better explained by drift-diffusion. In contrast, for the remaining 31 cells that exhibit plateaus greater than 2 sp/s only 4 cells were classified as diffusion. This is consistent with our diagnosis: the diffusion model is less penalized by late spikes when there are fewer of them (i.e., when *α*_*out*_ is low), thus reducing the bias in favor of stepping.

One might expect that the bias attributed to the negative going rates would be mitigated in data obtained in a reaction time experiment, since these experiments do not contain long stretches of neural activity after the decision is formed. However, as we next show, the bias remains an important factor. Latimer et al. analyzed 16 neurons from the Shadlen lab [14], and concluded that stepping provided a superior account of the responses in 12 of the cases (Supplementary figure S25 of *the study*). We first replicated their analysis using their code (Figure 5, upper left graph in each box). We reasoned, however, that the negative going rates were again problematic and wondered if late spikes were once again the culprit. To test this, we applied a transformation to the spike trains that removes late spikes while preserving the time course of the firing rates, as follows. We fit the stepping model to obtain the mean level of firing rate associated with a transition to the low-rate state (*α*_*out*_). For each trial, we reduced the number of spikes in each 50 ms bin by a random number drawn from a Poisson process with an expectation of *λ* = 0.05*α*_*out*_. The spikes that were removed were chosen at random within the 50 ms bin. The procedure achieves a reduction in the firing rate while preserving the time course of the firing rates in the averages as well as the detailed dynamics of the spike trains on single trials (right graph in each box).

**Figure 5.**
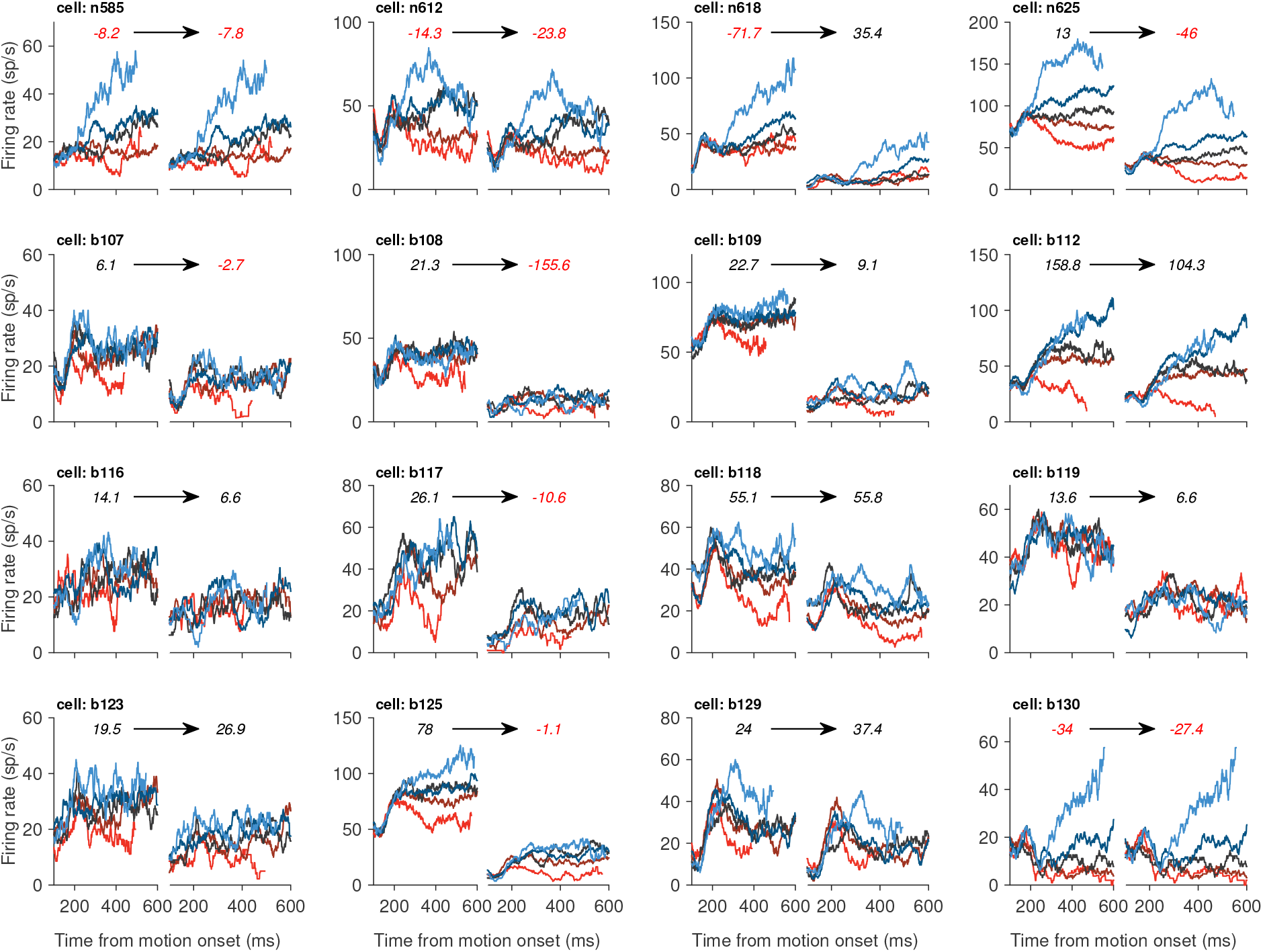
Reducing the overall spike rate of LIP neurons lessens the bias in favor of stepping. Each panel shows the firing rate from a different neuron (identified at the top of each panel). Within each panel, the left plot shows that rates obtained from the actual LIP data, and the right plots shows the rates after reducing the overall firing rate with the method described in the text. The numbers inside each plot indicate the ΔDIC. Positive (negative) numbers indicate support for stepping (diffusion) and are shown in black (red). The color of the solid lines denotes motion coherence, as in the previous figure.

We then applied the model comparison to this modified dataset. Five neurons that were originally classified as stepping were reclassified as diffusion; one neuron switched from diffusion to stepping. Overall, the reduction in firing rates doubled the number of neurons that were best accounted for by the diffusion model. This shows that the problem of unstopped negative drift rates is not limited to the data set from the Huk lab but is a factor in the analysis of data obtained in an RT experiment as well. As argued elsewhere, this is not the only source of bias in the model (see [12]).

## Discussion

Single neuron electrophysiology has furnished important insights about a variety of neural computations by characterizing the average firing rates over repeated presentations of stimuli, movements or task. The implicit assumption is that the firing rate average from one neuron derived from many trials is a proxy for the firing rate average of many neurons on one trial. The approach has delivered much of what we know about neural computation in cortex, where there is empirical support for the existence of many neurons that respond similarly. However, the drawbacks of this assumption are also well known (e.g., [17, 18]), hence an attempt to characterize a latent rate process for single trials is laudable. We congratulate Latimer et al. for attempting such a feat, but there are pitfalls, as we have shown.

The search for a latent rate in individual trials must be guided by a model of some kind, and any model comparison is limited by the quality of the models and data used to evaluate them. Here, we have focused on one particular problem in the instantiation of drift-diffusion. It offers an interesting cautionary tale because the model passes a validation test precisely because it cannot generate the very feature of the data that penalizes it—namely late spikes when the evidence is negative. This feature of the data is easily accommodated by stepping, thus biasing the model comparison, but it does not bear on the core issue of what the neurons do during decision formation. There is, after all, no controversy over the capacity of LIP neurons to sustain persistent activity at a high or low rate. The questions is what these neurons do as the decision is being formed. The view that appears most consistent with a variety of observations is that the firing rate undergoes a biased random walk—idealized as drift-diffusion—representing the accumulation of noisy samples of evidence. When the process stops, the persistent activity represents an oculomotor plan. In another brain area it might represent a plan to do something else, such as reach [9, 19] or implement a rule [20], perhaps.

The diffusion model employed in *the study* made a number of questionable assumptions, which were discussed in a brief Technical Comment [12]. The one we focused on here is the absence of a mechanism to stop negative going rates. This feature of the model makes it unable to account for late spikes on trials with strong negative motion. Whereas stepping naturally accommodates these late spikes, their diffusion model must either suffer a penalty for these spikes or adjust the drift rates erroneously (Figs 1 and 4). This is especially problematic in tasks in which stimulus duration is controlled by the experimenter—like the main task analyzed by Latimer et al.—because the stimulus duration can be much longer than the decision time. For example, for easy decisions (i.e., strong motion), decision time is on the order of 50-100 ms, which is but a small fraction of the 500-1000 ms duration of the spike trains analyzed in *the study*. This implies that there are trials with up to 950 ms of spiking activity that unfairly penalize the diffusion model for its lack of a mechanism to freeze the neuronal activity at a low positive rate after a decision has been made.

We showed that this deficiency biases the model comparison against diffusion. We did not remedy the model but added or removed spikes from data and simulations. A proper remedy would augment the diffusion model with a mechanism that allows the diffusion to stop at non-zero firing rates. However, implementing such a mechanism is not trivial, as it requires the modeler to make assumptions about what it is that stops these negative going processes. We concur with Latimer et al. that the responses leading to T_out_ and T_in_ choices are not mirror replicas. Presumably, the negative-going rates are stopped when a population of neurons that accumulate evidence in favor of the opposite target reach a bound [8, 21-23], and these response levels appear to be less stereotyped [8, 14].

One might be tempted to dismiss the comparison of diffusion and stepping as unimportant, because both mechanisms could support population dynamics that would implement the bounded accumulation of noisy evidence. This position would be mistaken, in our view, as it renders an understanding of neural mechanism a mere appendix to computational and psychological concept. We are thus aligned with Latimer et al. on this point. The central question raised by these authors is important and fundamental. Do the responses of single neurons like those in LIP represent graded quantities, or are these gradations of firing rate represented by the average over a population of neurons, each capable of emitting only a few stable firing rates? We think the best way to address this question is experimentally [24], but there is no doubt that sophisticated analyses of data, using techniques like those in *the study,* will contribute to this effort.

## Acknowledgments

This research was supported by the Howard Hughes Medical Institute and the National Eye Institute (R01 EY11378). We thank Shin Kira and Daniel Wolpert for helpful discussions, and Chris Fetsch, Yul Kang, Danique Jeurissen, Josh Gold and Bill Newsome for comments on the manuscript.

